# Sex-Related Neurocircuitry Supporting Camouflaging in Adults with Autism: Female Protection Insights

**DOI:** 10.1101/2021.11.03.466990

**Authors:** Melissa J. M. Walsh, Broc Pagni, Leanna Monahan, Shanna Delaney, Christopher J. Smith, Leslie Baxter, B. Blair Braden

**Author notes:** Correspondence to: B. Blair Braden, Melissa J. M. Walsh, Full address: College of Health Solutions, Arizona State University, 975 S. Myrtle Ave., Tempe, AZ 85281, USA, E-mail.

## Abstract

The male preponderance in autism led to the hypothesis that aspects of female biology are protective against autism. Females with autism report engaging in more compensatory behaviors (i.e., “camouflaging”) to overcome autism-related social differences, which may be a downstream result of protective pathways. No studies have examined sex-related brain pathways supporting camouflaging in females with autism, despite its potential to inform mechanisms underlying the sex bias in autism.

This case-control design included 45 non-intellectually-disabled adults with autism (male/female: 21/24) and 40 neurotypical adults (male/female: 19/21) ages 18-71. We used group multivariate voxel pattern analysis to conduct a data-driven, connectome-wide characterization of “sex-atypical” (sex-by-diagnosis) and “sex-typical” (sex) brain functional connectivity features linked to camouflaging, and validated findings in females with autism multi-modally via structural connectometry. Exploratory associations with cognitive control, memory, emotion recognition, and depression/anxiety examined the adaptive nature of functional connectivity patterns supporting camouflaging in females with autism.

We found 1) “sex-atypical” functional connectivity patterns predicting camouflaging in the hypothalamus and precuneus and 2) “sex-typical” patterns in the anterior cingulate and right anterior parahippocampus. Higher hypothalamic functional connectivity with a limbic reward cluster was the strongest predictor of camouflaging in females with autism (a “sex-atypical” pattern), and also predicted better cognitive control/emotion recognition. Structural connectometry validated functional connectivity results with consistent brain pathways/effect patterns implicated across multi-modal findings in females with autism.

This data-driven, connectome-wide characterization of “sex-atypical” and “sex-typical” brain connectivity features supporting compensatory social behavior in autism suggests hormones may play a role in the autism sex bias. Furthermore, both “male-typical” and “female-typical” brain connectivity patterns are implicated in camouflaging in females with autism in circuits associated with reward, emotion, and memory processing. “Sex-atypical” results are consistent with the fetal steroidogenic hypothesis, which would result in masculinized brain features in females with autism. However, female genetics/biology may contribute to “female-typical” patterns implicated in camouflaging.

## Introduction

Autism spectrum disorder (ASD) has an estimated sex bias of 3-4:1 males to females.^1,2^ This male preponderance suggests sex-related biology may protect females or increase male vulnerability.^3^ Accumulating evidence supports the female protection hypothesis, with diagnosed females showing greater ASD genetic liability than males.^4^ Sex/gender-related ASD models have been proposed to identify mechanisms underlying the sex bias. The Extreme Male Brain model presents evidence that ASD represents the extreme end of cognitive/behavioral masculinization,^5^ while the Gender Incoherence model highlights masculinized phenotypic qualities in females with ASD (ASD-F) and feminized qualities in males with ASD (ASD-M).^6^ Both models suggest ASD-F present with a more masculine phenotype. However, certain traits linked to core social symptoms are more “female-typical” in ASD-F (e.g., social attention, linguistic discourse skills, social motivation).^7^ Both “masculine” and “feminine” sex-related biology may be implicated in the ASD sex bias, but characterization of relevant sex-related neurobiological pathways is understudied.

Most ASD risk genes are not sex-specific, and sex-related biology is thought to act downstream of risk variants to promote female protection.^8^ Transcriptomic studies show immune/microglial-associated genes are male-biased in neurotypicals (NT) and upregulated in ASD, while genes associated with neuronal functioning are female-biased in NT and downregulated in ASD,^8,9^ again implicating both masculine and feminine sex-related biology in ASD risk. To date, transcriptomic studies have examined bulk tissue from prefrontal, temporal, and cerebellar regions in small samples, and no studies have investigated brain regions showing pronounced sex differences (e.g., hypothalamus).^10^ Thus, inference about brain regions implicated in the ASD sex bias remains limited. Neuroimaging is a complementary approach to transcriptomics, allowing for *in vivo*, whole-brain analyses on larger, more homogeneous samples. Resting-state functional MRI (rs-fMRI) and diffusion tensor imaging (DTI) can be applied to characterize functional and structural pathways implicated in the ASD sex bias.^11^

A recent systematic review highlights that ASD-F show more atypical brain features than ASD-M, but some atypical features may be “protective.”^12^ For example, the left anterior prefrontal cortex has shown more atypical features in ASD-F.^13^ However, left anterior prefrontal functional connectivity (FC) with reward circuitry may buffer against ASD genetic risk in females.^14^ A novel way to elucidate brain systems underlying the sex bias in ASD is to characterize sex-related brain correlates of behaviors that show a sex bias in ASD. For example, greater use of compensatory strategies is reported in ASD-F than ASD-M,^15,16^ and this may be one mechanism by which ASD-F “work around” ASD genetic liability. The primary compensatory construct investigated in ASD is “camouflaging” (e.g., behaviors to mask or overcome ASD-related social differences).^17^ The field is new, and the first studies quantified camouflaging as the discrepancy between observable versus “intrinsic” (e.g., selfreport) ASD traits.^18,19^ The discrepancy approach has linked greater “camouflaging” in ASD-F to reduced cerebellar/medial temporal lobe volumes^18^ and increased activation of the medial prefrontal cortex during self-reflection^19^. However, discrepancy scores lack psychometric validation and may not accurately quantify camouflaging. More recently, the *Camouflaging Autistic Traits Questionnaire* (*CAT-Q*) was developed,^20^ but has yet to be linked with neuroimaging. Furthermore, no studies have examined brain connectivity camouflaging correlates, despite their translational utility.^21^ A recent neuroimaging genetics study found reward circuit FC within the left anterior prefrontal cortex was distinctly linked to greater genetic risk but reduced socio-cognitive symptom severity in ASD-F but not ASD-M.^14^ Thus, reward circuits may be candidate pathways supporting female protection, but the study’s seed-based methodology limits connectome-wide inference. Another recent study examined associations between FC and polygenic ASD risk, finding that greater genetic risk was linked to higher salience network FC with somatosensory cortex in boys than girls, irrespective of ASD diagnosis.^22^ Again, this study also used seed-based methodology, limiting connectome-wide inference. Data-driven approaches are needed to advance understanding of pathways implicated in the sex bias in ASD.

This study sought to examine sex differences in the neurocircuitry supporting camouflaging in ASD using a data-driven, connectome-wide approach. The objectives were to examine “sex-atypical” and “sex-typical” FC patterns predicting camouflaging in ASD. To characterize the adaptive nature of FC patterns linked to camouflaging in ASD, we examined associations with cognitive control, memory, emotion recognition, and depression/anxiety. Finally, we validated FC findings multi-modally using structural connectivity.

## Materials and methods

### Participants

The sample was derived from a larger age- and IQ-matched study (*n*=200) examining sex differences in non-intellectually-disabled, young-to-older adults with ASD. The study began in 2015 and the *CAT-Q* was added in 2019. Participants were selected if they had complete *CAT-Q*, rs-fMRI, and DTI data, with a total of 85 participants ages 18-70 (Table 1; *n*=24 ASD-F, *n*=21 ASD-M, *n*=20 NT-F, *n*=19 NT-M). As no prior studies have investigated sex-related functional connectivity patterns predicting compensatory behavior in ASD, adequate sample size was determined from a recent study examining sex-related circuits predicting genetic risk in ASD.^22^ With a slightly larger groupwise sample than our study (e.g., *n*=31 ASD-F compared to present study with *n*=24 ASD-F), this study reported modest peak-cluster effect sizes (*Z*~4),^22^ suggesting that our smaller sample will be adequate to detect sex-related brain-behavior associations

**Table 1.** Sample descriptive statistics and group differences.

|  | ASD |  | NT |  | Difference? |
| --- | --- | --- | --- | --- | --- |
|  | Females | Males | Females | Males |  |
| n | 24 | 21 | 21 | 19 |  |
| Age | 39.42<br>(13.21)<br>19-60 | 41.95 (11.39)<br>26-64 | 43.57<br>(15.68)<br>18-65 | 48.53<br>(13.35)<br>26-70 | None |
| CAT-Q | 120.21<br>(30.34)<br>49-164 | 107.00<br>(23.29)<br>64-161 | 63.95<br>(24.45)<br>38-139 | 72.68<br>(13.34)<br>55-101 | <sup>a</sup>ASD>NT,<br><sup>b</sup>ASD-F > ASD-M |
| IQ | 110.25<br>(16.94)<br>73-148 | 104.14<br>(15.95)<br>70-131 | 108.05<br>(12.71)<br>77-132 | 107 (11.24)<br>92-127 | none |
| SRS-2 | 103.87<br>(29.56)<br>44-167 | 110.43<br>(35.45)<br>21-155 | 21.95 (9.67)<br>5-46 | 26.95<br>(14.45)<br>7-57 | <sup>c</sup>ASD>NT |
| ADOS-2 | 9.87 (2.13)<br>7-14 | 10.67 (3.73)<br>7-19 | n/a | n/a | none |
| rs-fMRI Motion<sup>d</sup> | 0.06 (0.02)<br>0.02-0.08 | 0.07 (0.02)<br>0.02-0.11 | 0.06 (0.01)<br>0.04-0.09 | 0.07 (0.03)<br>0.02-0.13 | none |
| DTI Motion<sup>e f</sup> | 0.34 (0.13)<br>0.16-0.61 | 0.65 (0.61)<br>0.19-3.01 | 0.31 (0.15)<br>0.12-0.64 | 0.60 (0.38)<br>0.30-1.63 | <sup>g</sup>male>female |

Differences were tested via ANOVA for main effects of diagnosis, sex, or their interaction. If significant, post-hoc t-tests examined pairwise group differences driving the significant main effect or interaction. \*Camouflaging Autistic Traits Questionnaire (CAT-Q); Social Responsiveness Scale - 2nd Edition (SRS-2); <sup>a</sup>t<sub>83</sub>=8.62, p<.001; <sup>b</sup>t<sub>43</sub>=1.62, p=.11, <sup>c</sup>t<sub>83</sub>=15.25, p<.001; <sup>d</sup>CONN average motion observed (disregarding outlier scans; threshold = 0.5); <sup>e</sup>average root mean square displacements relative first volume; <sup>f</sup>no group differences in percent outlier slices or framewise displacement; <sup>g</sup>t<sub>83</sub>=3.79, p<.001

This sample has been characterized previously.^23,24^ In brief, ASD participants were recruited using the *Southwest Autism Research and Resource Center* lifetime database of voluntarily enrolled individuals who consented to be contacted for future studies and presentations at local ASD community events. For all participants, snowball recruitment and community fliers were used. ASD diagnoses were confirmed via the *Autism Diagnostic Observation Schedule-2*^25^ module 4, a brief case history, and the DSM-V checklist combined with clinical judgment. NT participants were enrolled if they had 1) no suspected diagnosis of ASD, 2) *Social Responsiveness Scale – 2^nd^ Edition*^26^ (*SRS-2*) T-scores ≤ 66, and 3) no first-degree family history of ASD. All participants were subject to the following exclusion criteria: 1) *Kaufman Brief Intelligence Test – 2^nd^ Edition* ^27^ scores <70, 2) *Mini Mental State Exam* ^28^ scores <26, 3) a history of head injury with loss of consciousness or neurological disorders, or 4) current seizure disorder or use of seizure medications. Psychiatric co-morbid conditions were not exclusionary given high prevalence in ASD^29^ (exception: schizophrenia). This study was conducted in compliance with Arizona State University’s ethical research standards and the Declaration of Helsinki 2000 revision. Participants provided Institutional Review Board approved written consent.

### Behavioral Measures

#### Camouflaging

The *CAT-Q*, published in 2018,^20^ is a self-report questionnaire developed through interviews of the “camouflaging” experience in adults with ASD. This questionnaire quantifies behaviors used to compensate for or mask autistic traits during social interactions, scored on a 7-point Likert scale from “Strongly Disagree” to “Strongly Agree.” Sample questions include “I have tried to improve my understanding of social skills by watching other people” and “I monitor my body language or facial expressions so that I appear interested by the person I am interacting with.” The total score was used in analyses. Given that camouflaging and ASD traits are correlated,^30^ the *SRS-2*^26^ measured self-reported ASD traits and was used as a covariate in all analyses to account for severity-related brain differences.

#### Neuropsychological Tests

Selected measures spanned broad executive function, memory, emotional, and psychiatric domains previously implicated in camouflaging or female protection. ^7,16,30–33^ In brief, the *Wisconsin Card Sorting Task,^34^ Stroop Color and Word Test*,^35^ and *Tower of London*^36^ estimated flexibility, cognitive control, and planning, respectively. The *Wechsler Memory Scale – Visual Reproduction II*^37^ and *Rey Auditory Verbal Learning Task*^38^ measured visual and verbal memory. The *Toronto Alexithymia Scale*^39^ and *Reading the Mind in the Eyes Task*^40^ measured emotional self-awareness and facial emotion recognition. Finally, the *State Trait Anxiety Inventory*^41^ and *Beck Depression Inventory-II*^42^ measured trait anxiety and depression. See Supplementary Methods S1 for detailed measure description.

### MRI Parameters

#### Acquisition

A 3T Philips Ingenia scanner collected anatomical, rs-fMRI, and DTI images (max. gradient strength=5 m T/m). The anatomical sequence was 3D magnetization prepared rapid acquisition gradient echo (MPRAGE; 170 axial slices, 1.2mm slices, 240mm FOV, 256×256 acquisition matrix). Six-minute (eyes closed) rs-fMRI scans were collected via a whole-brain coverage, gradient-echo, echo-planar (EPI) series (3000ms TR, 25ms TE, 80° flip angle, 3mm slices, 240 mm FOV, 64×64 acquisition matrix). DTI scans were acquired via a whole-brain coverage, diffusion inversion recovery sequence (7055.39ms TR, 118.655ms TE, 32 directions, 2500 ms/mm^2^ b-value, 1.41mm in-plane resolution, 3mm slice thickness).

#### Preprocessing and Subject-Level Connectivity Maps

Raw rs-fMRI images were first despiked (*Wavelet Despiking Toolbox*).^43^ Subsequent preprocessing was conducted in *SPM-12* (**https://www.fil.ion.ucl.ac.uk/**), including 1) functional image realignment, 2) anatomical image segmentation, skull-stripping, functional image co-registration, and 3) DARTEL normalization to MNI space. The CONN Toolbox identified functional outliers at a conservative 0.5mm framewise displacement threshold, followed by aCompCor confound regression,^44^ including realignment parameters/first order derivatives, scrubbing, linear detrending, and bandpass filtering [.008 .1]. FC maps were calculated using group multivariate voxel pattern analysis (MVPA). In brief, this procedure applied principle components analysis to each subject’s voxel-to-voxel correlation matrices to retain 64 components,^45^ then group principle components analysis to identify the top 15 salient spatial FC components across all subjects.

*FSL (FMRIB* software library, **www.fmri.ox.ac.uk/fsl**) was used for diffusion image preprocessing and *DSI Studio* (**www.dsi-studio.labsolver.org**) to estimate structural connectivity. In *FSL*, eddy correction was applied to account for head motion and eddy current distortions. Average root-mean-square displacement from the first non-diffusion weighted volume, scan-to-scan displacement, and percentage of outlier slices estimated subject-level motion (see Table 1 for group comparison). Eddy-corrected diffusion data was entered into *DSI Studio* to estimate quantitative anisotropy (QA). Unlike the more commonly estimated fractional anisotropy, which is thought to reflect the rate of diffusion, QA estimates the density of diffusing water in the fibers’ standard axonal direction as defined by an atlas.^46^ This procedure applied q-space diffeomorphic reconstruction (QSDR)^47^ to a standard template to calculate the spin distribution function^48^ with a 1.25mm diffusion sampling length and 1mm isotropic resolution. Using the *Human Connectome Project 1065* atlas, the spin distribution function magnitude in the standard fiber direction estimated voxel-wise QA, and values were concatenated within and across subjects to generate a local connectome matrix (Supplementary Methods S2 for detailed rs-fMRI and DTI preprocessing methods/connectivity map construction).

#### Group-Level Analysis

Using the CONN Toolbox, a general linear model was specified including group-wise intercepts and slopes to predict *CAT-Q* scores with age/*SRS-2* covariates (accounting for age/severity-related brain variability). Direction-agnostic group-MVPA contrasts tested positive/negative effects simultaneously. “Sex-atypical” patterns in ASD were modeled as inverse slopes for ASD-F/NT-M vs. ASD-M/NT-F. “Sex-typical” patterns were modeled as inverse slopes for females vs. males (See Supplementary Methods S3 for contrast details). An F-test evaluated all 15 group-MVPA rs-fMRI components simultaneously, but separately for each contrast. Post-hoc seed-to-voxel analyses characterized FC patterns driving the significant group-MVPA effects with positive/negative contrasts tested separately. For all rs-fMRI analyses, a voxel height threshold of p<0.001, uncorrected and cluster threshold of p<0.05, false discovery rate (FDR) corrected was used. To determine if results were driven by subgroups, we tested regression model significance using mean FC values from significant post-hoc clusters to predict residualized CAT-Q scores (partialing out age/*SRS-2*), with one model per MVPA cluster. Unthresholded t-maps from each MVPA post-hoc contrast were submitted to the Neurosynth Decoder (**https://neurosynth.org/decode/**) and word clouds were generated using R Studio Wordcloud2 for the top 10 correlated terms (based on correlation absolute values).

Group connectometry was conducted in DSI Studio^49^ using a whole-brain regression procedure, separately across each sex-by-diagnosis group, to identify tracts significantly associated with camouflaging. Given that ASD-F show more camouflaging than ASD-M^15^ and this may be a mechanism of female protection, structural connectivity correlates of camouflaging in ASD-M and NT groups were not of primary interest but are reported in Supplementary Figure S2. The model included CAT-Q as the study variable of interest, with age, SRS-2, and average root-mean-square displacement as nuisance covariates to account for age, symptom severity, and motion-related signal variability (given male>female group differences on this parameter; Table 1). A t-threshold of 3, corresponding to the more conservative recommended threshold in *DSI Studio*, was used to select QA values associated with the study variable for subsequent deterministic fiber tracking along substantial coefficients.^50^ Topology-informed pruning^51^ was applied across four iterations to filter false tracks. For tracks exceeding a length of 40 voxels, FDR<.05 was estimated using a distribution of 4000 random permutations to subject phenotypic values. Average track QA was plotted against residualized CAT-Q scores (age/SRS-2 partialed out) to visualize effects.

#### Functional Connectivity-Behavior Associations

For ASD-F, two-tailed partial Pearson correlations (age/*SRS-2* covariates) were calculated between behavioral measures and each FDR-corrected cluster from MVPA post-hoc results. Again, while associations in ASD-F were of primary interest given higher self-reported use of camouflaging,^15^ we report behavioral associations in ASD-M in Supplementary Fig. S3.

#### “Sex-Agnostic” (Diagnosis) Effects

Diagnosis-differential (“sex-agnostic”) camouflaging-FC (Supplementary Fig. S1) and behavioral associations (Supplementary Fig. S4) were explored. Given the purpose of this paper to characterize sex-related circuits underpinning camouflaging in ASD, the Methods/Results/Discussion are documented in Supplementary Materials.

## Results

### Functional Connectivity

#### “Sex-Atypical” (Sex-by-Diagnosis) Effects

Group-MVPA revealed clusters in the precuneus and hypothalamus (Fig. 1; Table 2). For both regions, ASD-F showed “male-typical” camouflaging-FC associations. ASD-M showed inverse “female-typical” associations that were non-significant for precuneus but significant for hypothalamus FC. For the precuneus, greater FC with the left dorsolateral prefrontal cortex and bilateral superolateral occipital cortex (Supplementary Table S2) predicted less camouflaging in ASD-F/NT-M and inverse patterns in ASD-M/NT-F. For the hypothalamus, greater FC with a left lateralized limbic cluster (basal forebrain/left orbitofrontal cortex, left ventral striatum/thalamus/substantia nigra) and the right anterior cingulate cortex as well as lower FC with a cerebellar cluster (bilateral crus 2/1, right lobule 6) predicted higher levels of camouflaging in ASD-F/NT-M and lower in ASD-M/NT-F.

**Figure 1.**
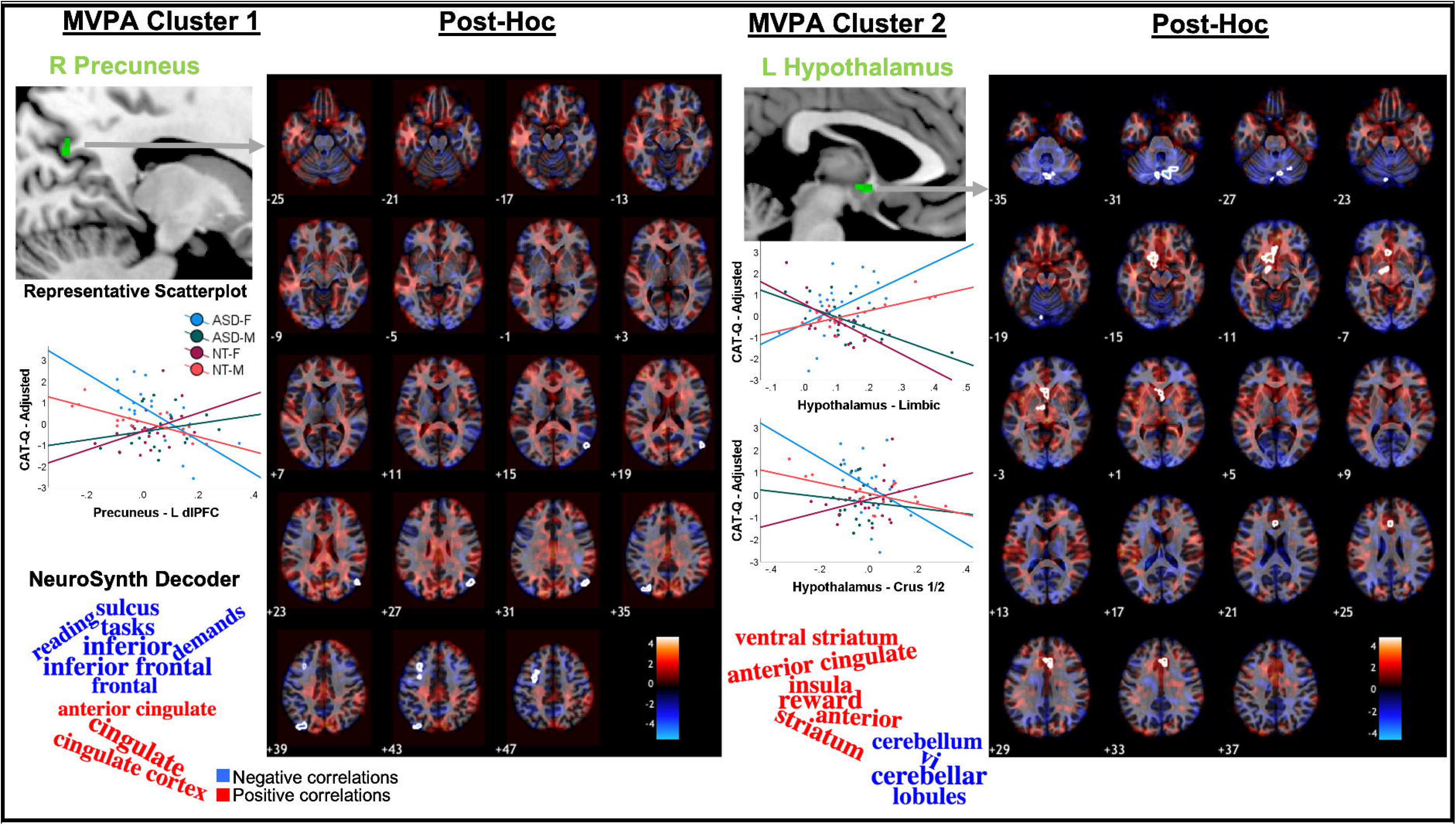
MVPA-derived clusters showing “sex-atypical” FC patterns predicting camouflaging in ASD. ASD-F showed “male-typical” and ASD-M showed “female-typical” FC-camouflaging associations. Scatterplots display group-wise linear effects for the most significant clusters surviving FDR-correction for both positive and negative contrasts (if significant) plotted against CAT-Q scores (adjusted to account for age and severity-related variability). Word clouds were generated by submitting unthresholded t-maps from post-hoc MVPA seed-to-voxel contrast to the Neurosynth Decoder and plotting the top 10 terms (based on correlation absolute values). *right (R); left (L)

**Table 2.**
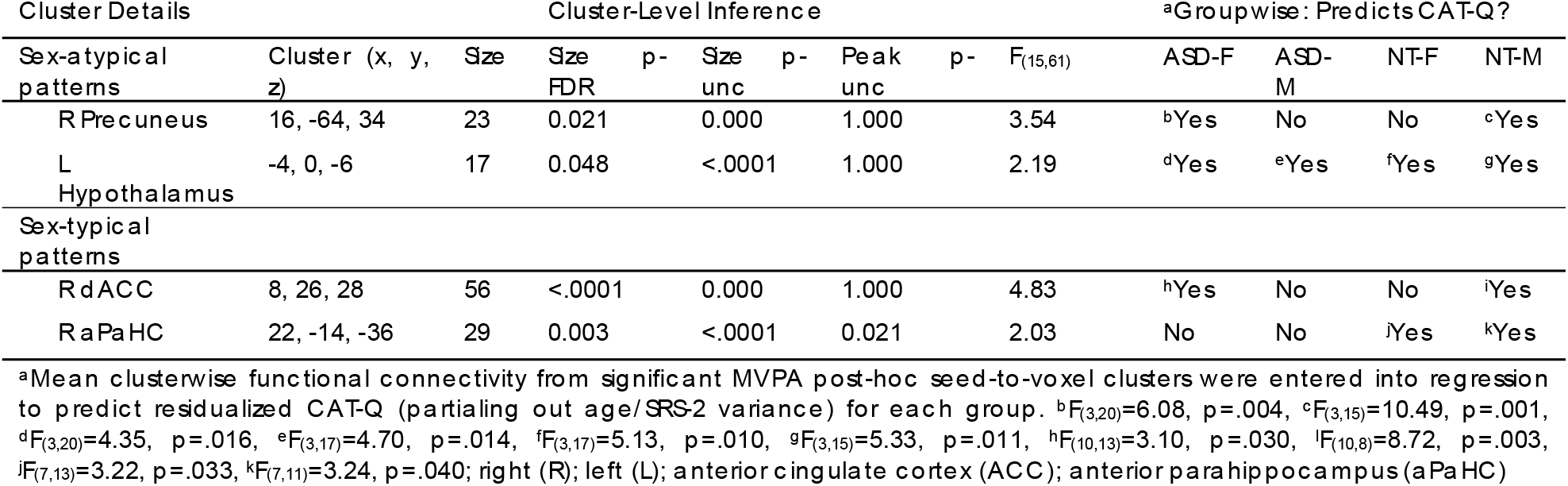
Sex-atypical and sex-typical camouflaging-functional connectivity associations in ASD revealed via group-MVPA.

#### “Sex-Typical” (Sex) Effects

This contrast revealed significant MVPA clusters in the right anterior cingulate and right anterior parahippocampus (aPaHC; Fig. 2; Table 2). Greater right anterior cingulate FC with bilateral sensorimotor, lateral parietal, thalamic, and lingual regions; right ventral prefrontal cortex and anterior temporal cortex; left dorsolateral prefrontal cortex; and mid-cingulate cortex (Supplementary Table S2) predicted more camouflaging in females and less in males. For the right anterior parahippocampus, higher FC with bilateral parietal and left ventral prefrontal cortex as well as lower FC with the right temporal pole predicted more camouflaging in females and less in males.

**Figure 2.**
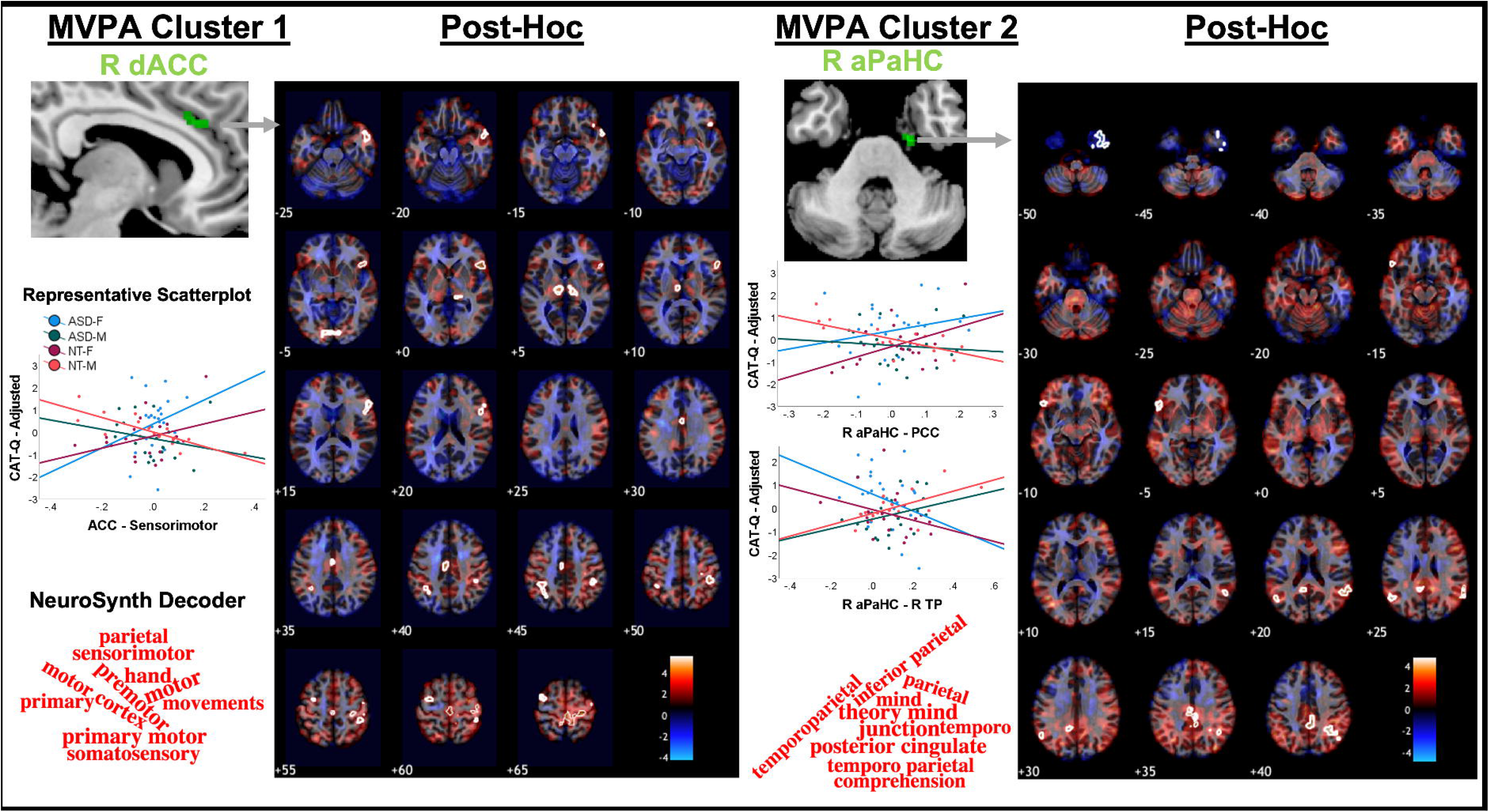
MVPA-derived clusters showing “sex-typical” FC patterns predicting camouflaging in ASD. ASD-F showed “female-typical” and ASD-M showed “male-typical” camouflaging-FC associations. Scatterplots display group-wise linear effects for the most significant clusters surviving FDR-correction for both positive and negative contrasts (if significant) plotted against CAT-Q scores (adjusted to account for age and severity-related variability). Word clouds were generated by submitting unthresholded t-maps from post-hoc MVPA seed-to-voxel contrast to the Neurosynth Decoder and plotting the top 10 terms (based on correlation absolute values). *right (R); dorsal anterior cingulate cortex (dACC); anterior parahippocampus

### Behavior Associations

FC patterns positively predicting camouflaging in ASD-F were generally linked to better executive functioning (Fig. 3). Intriguingly, hypothalamic-limbic FC was the strongest positive predictor of camouflaging and also predicted better cognitive control and facial emotion recognition. Specific to the right anterior cingulate, higher FC predicted greater camouflaging but poorer emotional self-awareness. FC patterns negatively predicting camouflaging showed a general pattern, albeit non-significant, where higher FC predicted poorer executive functioning.

**Figure 3.**
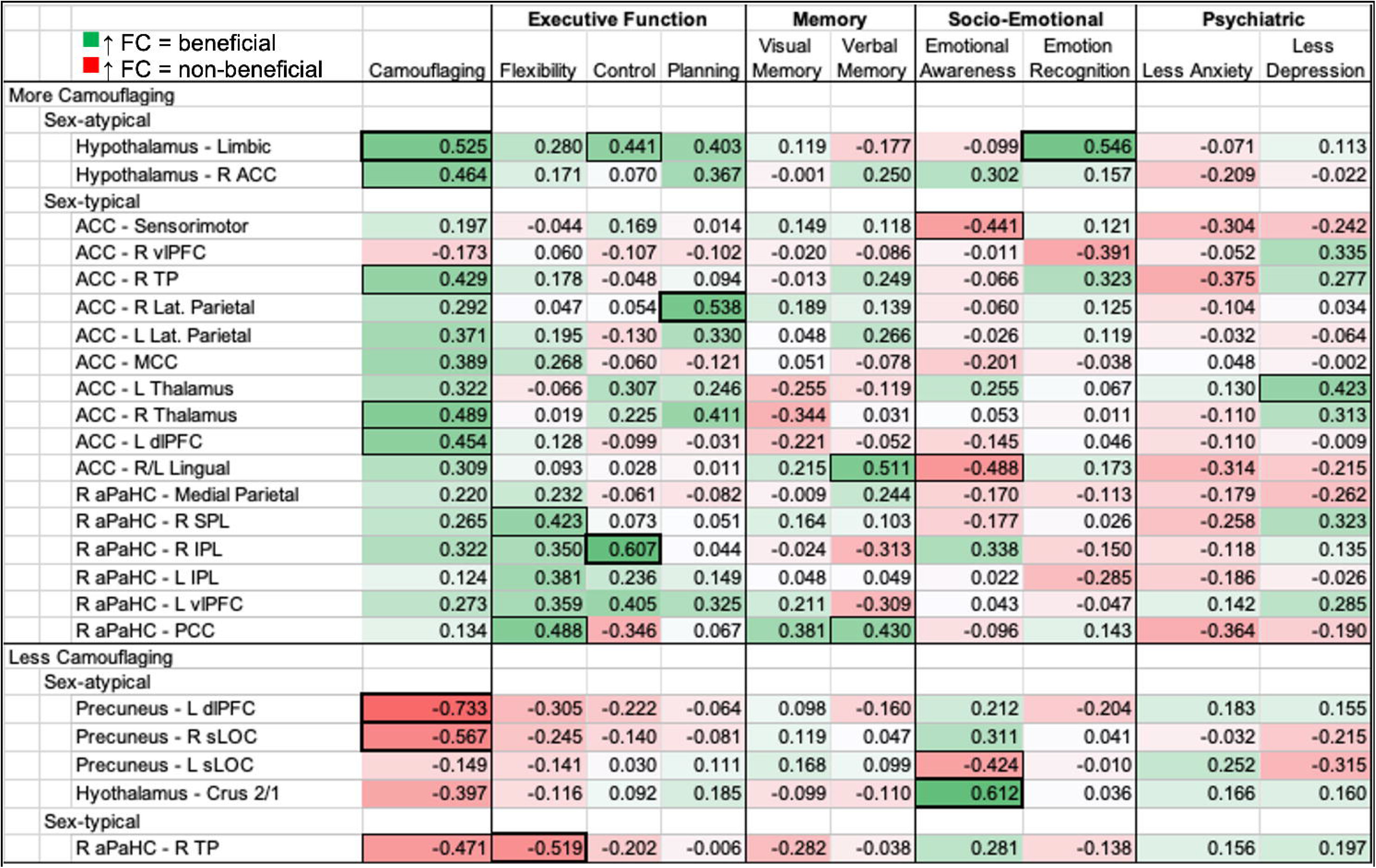
Exploratory behavioral associations in ASD-F to examine the adaptive nature of sex-related FC patterns implicated in camouflaging. Coefficients are partial Pearson correlations with SRS-2/age covariates. Dark bold boxes indicate p≤.01 and thin outlines p≤.05. For all subscales, higher correlations indicate higher FC is linked to better performance on cognitive/emotional tasks or attenuated anxiety/depression. Correlations were inverted (multiplied by −1) if higher scores on a cognitive/memory/psychiatric symptoms scale indicated worse performance/symptoms. *Camouflaging Autistic Traits Questionnaire Total Score (Camouflaging); Wisconsin Card Sorting Task Perseverative Errors (Flexibility); Stroop Color and Word Test Interference Score (Control); Tower of London Total Correct (Planning); Weschler Memory Test - Visual Reproduction Delayed Recall (Verbal Memory); Rey Auditory Verbal Learning Task A7 (Verbal Memory); Toronto Alexithymia Scale - 26 (Emotional Awareness); Reading the Mind in the Eyes Task (Emotion Recognition); State Trait Anxiety Inventory Trait Score (Anxiety); Beck Depression Inventory - II (Depression); Right (R); Left (L); Anterior Cingulate Cortex (ACC); ventrolateral prefrontal cortex (vIPFC);

FC patterns positively linked to camouflaging in ASD-F were negatively linked in ASD-M. For the anterior cingulate, higher FC predicted less camouflaging, poorer cognitive control, better emotional self-awareness (also for hypothalamic FC), and trends toward less depression/anxiety (Supplementary Fig. S3). For other FC patterns, associations were nonsignificant or mixed in positive and negative directions for executive functioning, memory, and facial emotion recognition. FC positively predicting camouflaging in ASD-M showed largely non-significant behavioral associations.

### Multi-Modal Validation: Structural Connectometry

#### Connectometry in ASD-F

QA was positively associated with the camouflaging variable (FDR<.05) in a track (total streamline count: 989) including the left anterior thalamic radiations, forceps minor, bilateral parahippocampal cingulum, and bilateral cerebellum/vermis (Fig. 4). Negative correlations were found in a track (streamlines: 1560) comprising posterior commissural fibers (corpus callosum [CC]-body, tapetum, forceps major), and the bilateral arcuate/superior longitudinal fasciculus (SLF).

**Figure 4.**
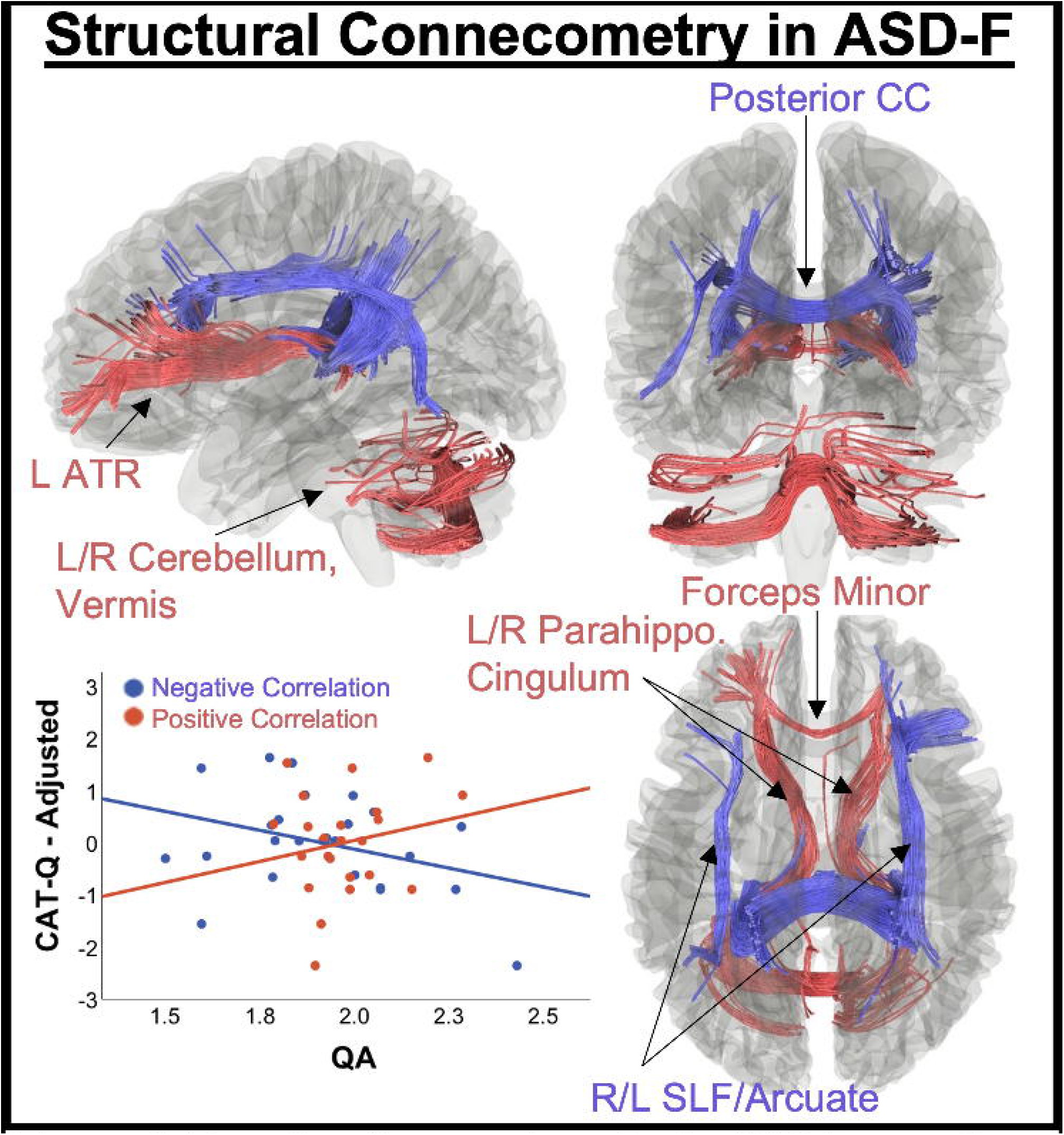
Results of connectometry in ASD-F revealing tracks whose structural connectivity (QA) correlated with camouflaging (p<.05, FDR-corrected). Positive correlations were found in a track comprising the left anterior thalamic radiation (ATR), forceps minor, bilateral parahippocampal cingulum, and bilateral cerebellum/vermis. Negative correlations were found in the posterior corpus callosum (body/tapetum/forceps major) and bilateral arcuate/superior longitudinal fasciculus (SLF). *age, SRS-2, and DTI motion (average displacement relative to 1^st^ non-diffusion volume) included as covariates in connectometry

#### Connectometry in ASD-M and NT Groups

In ASD-M, QA was negatively associated with camouflaging only in the cerebellum (right hemisphere/vermis; 870 streamlines; Supplementary Fig. S2). In NT groups, sparse connectometry associations were observed, mostly localized to cerebellar tracts. NT-F showed positive associations in the right cerebellum and CC-tapetum (streamlines: 35), and NT-M showed positive associations in the middle cerebellar peduncle and left frontal aslant tract (streamlines: 22).

## Discussion

Using a data-driven, connectome-wide analytic approach, we found distinct sex-related brain connectivity patterns supporting camouflaging in ASD-F. “Male-typical” FC patterns were found in the precuneus and hypothalamus and “female-typical” patterns in the right anterior cingulate and anterior parahippocampus. Hypothalamic-limbic FC significantly predicted camouflaging across each group (ASD-F, ASD-M, NT-F, NT-M), with “male-typical” positive associations in ASD-F and “female-typical” negative associations in ASD-M. This finding may suggest “gender incoherent”^6^ FC between the brain’s hormone center and reward circuitry contributes to sex-biased camouflaging patterns in ASD. In ASD-F, higher hypothalamic-reward circuit FC may be a particularly adaptive compensatory pattern given its 1) strong positive association with camouflaging, 2) its link to better cognitive control and facial emotion recognition, and 3) absent associations with depression/anxiety. Groupwise structural connectometry revealed substantial tracts associated with camouflaging in ASD-F, and sparse correlates outside of the cerebellum for ASD-M and NT groups. Furthermore, tracts implicated in camouflaging in ASD-F showed consistent correlation patterns to FC results and implicated overlapping circuits.

### Sex-Related FC Patterns Supporting Camouflaging

Higher hypothalamic FC with reward centers predicted greater camouflaging in ASD-F (“male-typical”) and less camouflaging in ASD-M (“female-typical”). In ASD-F, higher hypothalamic-limbic FC also predicted better cognitive control and facial emotion recognition. The hypothalamus is a sexually dimorphic structure,^52^ responsible for neuroendocrine regulation of the autonomic nervous system.^53^ Although high resolution spatial inference should be interpreted with caution, the hypothalamic region identified in this study overlapped most substantially with the paraventricular nucleus of the hypothalamus,^54^ implicated in sex-differential regulation of social behavior.^55^ Previous evidence has implicated reward centers in hormonally-mediated female protection in ASD, such that more ASD-associated oxytocin receptor allele variants uniquely predict greater left prefrontal-reward center FC in ASD-F, and this FC pattern is linked to reduced ASD socio-cognitive difficulties.^14^ Furthermore, reward centers play a role in sex-differential development of social behavior,^56^ motivated recruitment of cognitive control,^57^ and emotional learning.^58^ Speculatively, hypothalamic-reward center FC in ASD-F may support greater social motivation,^59^ facilitating cognitive control recruitment to improve social interactions and cue learning for accurate emotion recognition. The limbic reward cluster comprised dopaminergic regions implicated in reward processing (e.g., nigrostriatal pathway)^58^ and a hub of the cholinergic systems implicated in state arousal (e.g., basal forebrain).^60^ This may suggest “gender incoherent” sex steroid influence on major neurotransmitter systems in ASD contribute to sex-biased camouflaging behavior. Finally, greater hypothalamic-cerebellar FC was linked to less camouflaging in ASD-F (“male-typical”) and greater camouflaging in ASD-M (“female-typical”). This circuit plays a role in implicit socio-emotional processing^61^ and more intentional processes may be used by ASD-F.

The right precuneus also showed “sex-atypical” FC patterns supporting camouflaging in ASD. Greater FC with the left dorsolateral prefrontal cortex and bilateral superolateral occipital cortex predicted less camouflaging in ASD-F (“male-typical”) and inverse patterns in ASD-M (“female-typical”). The precuneus is a hub of the default mode network implicated in task-negative reflective states and dynamic switching with task-positive networks.^62^ Diverse term associations yielded by the Neurosynth Decoder may reflect the precuneus’ role in communication between task-positive and task-negative networks. A recent study found that lower estrogen/progesterone receptor expression in post-chemotherapy breast cancer patients predicted greater precuneus-dorsolateral prefrontal cortex FC and poorer cognitive functioning.^63^ Associations in ASD-F, while non-significant, trended toward greater FC predicting poorer cognitive control. A mechanism underlying the precuneus’s “sex-atypical” role in camouflaging in ASD may be atypical sex steroid receptor expression.

Right anterior cingulate FC with sensorimotor regions was a “sex-typical” FC pattern linked to camouflaging in ASD, such that higher FC predicted more camouflaging in females and less in males. The anterior cingulate is a hub of the salience network, involved in stimulus salience processing.^64^ Altered sensory processing is a hallmark of ASD,^65^ and greater FC between the anterior cingulate and sensorimotor regions may indicate attentional bias toward sensorimotor representation. Prior evidence shows that, irrespective of diagnosis, boys but not girls show a relationship such that increasing polygenic ASD risk predicts greater salience network FC with sensorimotor cortex.^22^ Thus, female biology may protect against ASD-risk gene mediated alterations in sensorimotor processing. Our data may further suggest that this more male penetrant feature of increased salience network-sensorimotor FC in ASD may reduce compensatory behaviors in ASD-M but increase them in ASD-F. We also found that greater anterior cingulate-sensorimotor FC predicted poorer emotional self-awareness in ASD-F but greater emotional self-awareness in ASD-M. The salience network plays a role in emotional and empathic processing,^66^ and FC with the sensorimotor cortex is implicated in somatic representation of emotions.^67,68^ The inverse associations between emotional awareness and anterior cingulate-sensorimotor FC in ASD-M vs. ASD-F (Supplementary Fig. S3) may be explained by sex hormones. For example, animal studies have shown that estrogen and testosterone inversely impact emotion context generalization.^69^ Similar sex-differential hormone influence on emotional learning may apply in ASD.

“Sex-typical” camouflaging-FC associations in ASD were also found in the right anterior parahippocampus. Higher FC with regions of the default mode network (e.g., precuneus) predicted more camouflaging in females and less in males. The parahippocampus acts as a hub linking the DMN with the medial temporal memory system^70^ and may process stimulus familiarity.^71^ Greater right parahippocampal-precuneus FC has been associated with better episodic remembering.^72^ This FC pattern may support familiarity encoding and autobiographical retrieval to improve social interactions in females. Accordingly, ASD-F showed positive associations between anterior parahippocampal FC and cognitive control/flexibility. From a neurochemical perspective, engagement of the right anterior parahippocampus/anterior cingulate and precuneus during memory tasks in women is altered by estrogen and anti-cholinergic drugs.^73^ These regions may constitute neuromodulatory hubs, where hormones and neurotransmitters interact to influence state arousal.

### Multi-Modal Validation

Structural connectivity associations with camouflaging were substantial in ASD-F, and largely absent outside of cerebellar tracts for ASD-M and NT groups. Furthermore, structural connectivity correlates of camouflaging in ASD-F broadly aligned with sex-related circuits identified in FC results. For example, higher structural connectivity in limbic and cerebellar tracts was linked to greater camouflaging in ASD-F. The left-dominant anterior thalamic radiations, anterior commissural fibers, and the medial cingulum form part of the limbic pathway connecting the hypothalamus with emotion, cognitive control, reward, and cerebellar circuits.^74^ These tracts largely paralleled “sex-atypical” FC results implicating hypothalamic FC with orbitofrontal/ventral striatal, anterior cingulate, and cerebellar regions. Furthermore, the direction of effects was largely consistent, where higher connectivity predicted more camouflaging in ASD-F. In addition, the cingulum was implicated in positive structural connectivity-camouflaging associations in ASD-F. This tract connects medial regions of the DMN^75^ and the parahippocampus,^76^ which is consistent with “sex-typical” results showing that higher right anterior parahippocampal FC with the DMN predicts more camouflaging in ASD-F. Finally, higher structural connectivity in parietal tracts connecting medial/lateral parietal and occipital cortex^77^ to precentral^78^ and dorsolateral prefrontal regions^79^ predicted less camouflaging in ASD-F. This finding is largely consistent with “sex-atypical” FC results in the right precuneus, where greater FC with superolateral occipital and dorsolateral prefrontal cortex predicted less camouflaging in ASD-F. The consistent direction of camouflaging-connectivity associations and coherence of circuits implicated across FC and structural connectivity analyses validates FC findings and suggests they are not influenced by scanner-related confound variables (e.g., head motion).

### Sex-Related Biological Mechanisms

The clusters identified in MVPA analyses; including the hypothalamus, precuneus, right anterior cingulate, and anterior parahippocampus; are all sex-hormone sensitive regions^73,80^. Various lines of evidence have linked elevated prenatal sex steroid exposure^81–85^ to ASD. Prenatal sex steroid exposure in female mice results in “masculinized” brain features.^86^ Thus, such exposure would be expected to result in some brain features being more “male-typical” in ASD-F, in alignment with observed connectivity patterns in this study. However, sex steroid influence cannot be easily disentangled from other aspects of sex-related biology. For example, a recent study highlights the X-chromosome’s privileged influence on neuroanatomical variation.^87^ Regions linked to enriched X-chromosome morphological influence overlap with the “sex-typical” right anterior cingulate/anterior parahippocampus and the “sex-atypical” precuneus nodes linked to camouflaging in this study. It is plausible that processes like X inactivation/escape may preserve function in sex-related pathways affected by ASD risk genes.^4^ The results of this study suggest a role for hormonally-mediated processes in sex-biased compensatory behavior in ASD, but other aspects of sex-specific biology (e.g., sex chromosomes) may also play a role and require further investigation.

### Limitations

The small study sample and high model complexity is a limitation. However, the biological plausibility of findings revealed via data-driven methods is encouraging. To ensure results are not sample-specific, replication is needed. Furthermore, the sample age range was broad (18-70 years) and inclusion of pre-/post-menopausal women and younger/older men suggests diverse circulating sex hormone levels. However, larger hormonal variance may improve sensitivity in these dimensional analyses. Testing whether results extend to younger cohorts is an important future direction. Exploratory investigations of the adaptive nature of camouflaging FC patterns via correlations with behavior were uncorrected for multiple comparisons. Future investigations would benefit from larger samples with rich phenotypic data to validate the current results. Finally, it should be noted that this study is not advocating the need to compensate for/camouflage social differences in ASD. We acknowledge camouflaging is associated with poorer mental health in ASD.^16^ Instead, the purpose of this study was to capitalize on observed sex difference in self-reported camouflaging in ASD due to its potential sensitivity to neurobiological differences related to the sex bias.

### Conclusion

This study is the first to characterize sex-related connectivity correlates of compensatory social behaviors in ASD (e.g., camouflaging) using a data-driven, connectome-wide neuroimaging approach. This study lends insight into the ASD sex bias, suggesting both “male-typical” and “female-typical” neurobiological pathways play a role in camouflaging in ASD-F. Circuits liked to greater camouflaging in ASD-F were associated with reward (“sex-atypical” pathway), emotion, and memory retrieval (“sex-typical” pathways). Intriguingly, higher FC between the hypothalamus and limbic reward circuits predicted more camouflaging, better cognitive control/emotion recognition, and showed no association with depression/anxiety in ASD-F, suggesting this “male-typical” FC pattern may be particularly adaptive. The observed “male-typical” patterns supporting camouflaging in ASD-F are consistent with the fetal steroidogenic hypothesis,^81–85^ given that such exposure in females would result in “male-typical” brain features.^12^ However, the growing literature on sex differences in the brain suggests that both masculine and feminine processes interact with individual genetics and environment across development in a time-sensitive manner to produce an individual’s brain “mosaic.”^88–90^ In keeping, the “female-typical” FC patterns linked to camouflaging suggest that female sex chromosomes or reproductive biology (e.g., ovarian hormones) also influence social compensatory behavior in ASD-F and, potentially, the broader sex bias in ASD. Further research is warranted investigating sex-differential brain and behavioral development in ASD across the lifespan, especially during sensitive windows of hormone transition, and its relationship to sex-specific biology (e.g., hormones, genetics).

## Acknowledgements

We are grateful to our participants who made this study possible, and to Sharmeen Maze for her dedication to pristine MRI data collection.

## Abbreviations

ASD: autism spectrum disorder
NT: neurotypical
ASD-F: females with ASD
ASD-M: males with ASD
NT-F: NT females
NT-M: NT males
rs-fMRI: resting-state functional MRI
DTI: diffusion tensor imaging
FC: functional connectivity
CAT-Q: camouflaging autistic traits questionnaire
SRS-2: social responsiveness scale – 2^nd^ edition
MVPA: multivariate voxel pattern analysis
QA: quantitative anisotropy

