## Supplementary Materials for "Sex-Related Neurocircuitry Supporting Camouflaging in Adults with Autism: Female Protection Insights"

**Supplementary Methods**

1. **Description of Measures for Exploratory Behavioral Analyses**

**Cognition**: The *Stroop Color and Word Test*,^1^ *Wisconsin Card Sorting Task*,^2^ and *Tower of London*^3^ were selected to estimate cognitive control, cognitive flexibility, and planning, respectively. The *Stroop Color and Word Test*^1^ is thought to measure inhibition of cognitive interference. Participants complete three conditions: word reading in black ink (Stroop I), naming of color patches (Stroop II), and color-words printed in either color-congruent or color-incongruent ink (e.g., the word “red” written in blue; Stroop III). In the final condition, participants are instructed to name the color of the ink, thus requiring inhibition of word reading. For each condition, participants are given 45 seconds to read/name items and the total number of completed items is marked. Interference scores are calculated as the interference = Stroop III – [(Stroop I + Stroop II)/2].^4^ Lower (e.g., more negative) interference scores indicate greater inhibitory challenges. For the *Wisconsin Card Sorting Task*, participants are instructed to draw cards from a deck and match each card to one of four key cards, but no instructions are given regarding matching rules. Participants are given correct/incorrect feedback after they place each card and are instructed to try their best to get the next card correct. The administrator changes the matching rule at regular intervals without advising the participant. The perseverative errors score was selected to measure cognitive flexibility. Higher scores indicate greater difficulties with rule shifting. Finally, for the *Tower of London*, participants are presented with a pegboard containing colored beads in a target position and instructed to match the bead position on their board in as few moves as possible from a given start position. The Total Correct score was selected to measure planning accuracy, with higher scores indicating better planning performance.

**Socio-Emotional**: The *Reading the Mind in the Eyes Task*^5^ and *Toronto Alexithymia Scale*^6^ were selected to measure facial emotion recognition and emotional self-awareness, respectively. For the *Reading the Mind in the Eyes Task*, participants are presented with pictures of the eyes of male or female characters and are instructed to select, from a set of four options, the emotion that best represents how the character is feeling. The *Toronto Alexithymia Scale* is a self-report measure used to diagnose alexithymia, a condition that has been linked to ASD and is characterized by poor awareness of one’s own emotions^7^. Higher scores indicate poorer emotional self-awareness.

**Memory**: The *Rey Auditory Verbal Learning Task*^8^ Delayed Recall and *The Wechsler Memory Scale – Visual Reproduction*^9^ Delayed Recall scores were selected as measures of delayed verbal and visual memory, respectively. For the *AVLT*, participants are asked to attend to a list of 15 words and immediately repeat back as many words as they can remember. They are presented with the same list across five trials. They are then presented with a new list of 15 words and again, asked to repeat back as many words as they can remember. Then, they are asked to recall as many words as they can from the first list, and again after a 20 minute delay (delayed recall, trial A7). For the *WMS – VR* task, participants are presented with a geometric design for 10 seconds, then the design is covered from view, and the participant must immediately draw the design from memory. After an approximately 20-minute delay, the participant is then asked to reproduce the images from memory.

**Psychiatric Co-morbidities**: The *Beck Depression Inventory-II*^10^ and *State Trait Anxiety Inventory*^11^ were used to measure depression and trait anxiety, respectively. These are self-report measures with good psychometric properties in NT cohorts. Higher scores indicate more depression and anxiety.

2. **Neuroimaging Data Preprocessing and Connectivity Map Construction**

**rs-fMRI**: Each subject’s raw and preprocessed data underwent visual inspection for major scanning-related or preprocessing artifacts. Furthermore, accurate co-registration of the functional and skull-stripped structural images was verified in native space. After DARTEL normalization, accurate MNI boundary registration was verified. For data-driven, connectome-wide group-MVPA, PCA was conducted separately for each subject to retain the top 64 components explaining variance in their voxel-to-voxel correlation structure^12^. Subsequently, group-PCA was conducted across all subjects for each voxel, separately, to retain the top 15 most salient spatial components. These components represent the between-subject variance in each seed-to-voxel FC map, ordered according to variance explained. The top 15 components were selected to fulfill a 5:1 subjects-to-components ratio^13^, after accounting for the 10 degrees of freedom used in GLM modeling of groupwise intercepts, slopes, and covariates.

**DTI**: Eddy-corrected images were entered into *DSI Studio* to estimate quantitative anisotropy. A batch quality control procedure checked for consistent image dimensions, resolution, diffusion image count, and b-table accuracy^14^. The neighboring diffusion weighted image correlation was calculated for each scan across similar-gradient volumes^15^. Higher correlation coefficients suggest higher image quality, and outlier scans were defined as those falling three absolute deviations (no outliers detected). The local connectome fingerprint (LCF) was selected as the diffusion metric of interest due to its sensitivity to phenotypic traits^16^. Unlike fractional anisotropy which is thought to reflect the rate of diffusion, the LCF estimates the density of diffusing water in the fibers’ standard axonal direction as defined by an atlas (here-to-forth referred to as quantitative anisotropy; QA). To estimate QA, q-space diffeomorphic reconstruction (QSDR)^17^ is used to calculate the spin distribution function (SDF)^18^, with the SDF representing an orientation distribution function of the density of diffusing spins.

3. **Group-level Analysis Contrast Details**

**rs-fMRI**: “Sex-atypical” patterns in ASD were modeled as inverse slopes for ASD-F/NT-M vs. ASD-M/NT-F (e.g. given this order of intercepts/slopes [ASD-F, ASD-M, NT-F, NT-M]; intercepts: [0 0 0 0], slopes: [1 -1 -1 1] or [-1 1 1 -1]). “Sex-typical” patterns were modeled as inverse slopes for females vs. males (e.g., intercepts: [0 0 0 0], slopes: [1 -1 1 -1] or [-1 1 -1 1]).

In additional to primary sex-related analyses, we conducted exploratory examination of diagnosis-differential camouflaging-FC associations. “Sex-agnostic” (diagnosis) effects were modeled as inverse slopes for ASD vs. NT (e.g., intercepts: [0 0 0 0], slopes: [1 1 -1 -1] or [-1 -1 1 1]. Seed-to-voxel analyses characterized FC patterns driving the significant group-MVPA effects (positive and negative contrasts modeled separately)

**Supplementary Results**

1. **Sex-agnostic (Diagnosis) Effects**

**rs-fMRI**: In addition to “sex-typical” effects in the right aPaHC, diagnosis differential camouflaging-FC associations were observed in the right aPaHC (Supplementary Fig. S1 and Supplementary Table S1). Specifically, ASD groups showed greater FC linked to more camouflaging, and inverse associations were found in NT groups. This effect was driven by FC with bilateral parietal and right prefrontal regions (Supplementary Table S2), and FC patterns significantly predicted camouflaging only in NT groups (Supplementary Table S1). Similarly, diagnosis-differential camouflaging-FC associations were observed for the left temporal pole (TP), and these FC patterns significantly predicted camouflaging across all groups (Supplementary Table S1). This effect was driven by positive camouflaging-FC associations with right lateral parietal/occipital cortex and negative associations for left lateral and medial visual regions in ASD, but inverse patterns in NT.

**FC Associations with Behavior**: Higher FC was mostly linked to more camouflaging in ASD. In ASD-F, higher FC generally predicted better executive functioning. For ASD-M, higher FC generally predicted better planning but poorer memory. FC patterns linked to less camouflaging in ASD were found only for left temporal pole-visual cortex FC. However, behavioral associations were mixed for ASD-F and ASD-M (Supplementary Fig. S3).

**Supplementary Discussion**

1. **Integration of Sex-agnostic Results**

**Functional Connectivity**: The right aPaHC showed “sex-agnostic” FC patterns predicting greater camouflaging in ASD. Specifically, greater FC with regions implicated in working memory (e.g. intraparietal sulcus, dlPFC; Supplementary Fig. S1) was linked to more camouflaging in ASD and inverse NT patterns. The right aPaHC plays a role in encoding and maintenance of bound information in working memory^19^. Speculatively, this FC pattern may support binding new and old autobiographical information in memory to improve social interactions in ASD. This hypothesis is supported by positive associations between “sex-agnostic” aPaHC FC patterns and executive functioning in ASD-F and ASD-M (Supplementary Fig. S4). The left temporal pole (TP) was also implicated in sex-agnostic effects, such that lower FC with left lateral and medial visual cortex predicted more camouflaging. Left TP-visual FC is thought to underlie visual pattern recognition for object identification^20^. One study found the TP may have a top-down, modulatory effect on the ventral visual stream for integration of perceptual information during social tasks^21^.

**Supplementary Tables**

| **Supplementary Table S1.** "Sex-agnostic" (diagnosis-differential) camouflaging-functional connectivity associations in ASD revealed via group-MVPA. | | | | | | | | | | | |
| --- | --- | --- | --- | --- | --- | --- | --- | --- | --- | --- | --- |
|  | Cluster Details | | | Cluster-Level Inference | | | | ᵃPost-Hoc: Predicts CAT-Q? | | | |
|  |  | x, y, z | Size | Size p-FDR | Size p-unc | Peak p-unc | *F_(_*_15, 61)_ | ASD-F | ASD-M | NT-F | NT-M |
|  | R aPaHC, pTFusC | 28, -22, -36 | 21 | 0.019 | 0.000 | <.0001 | 5.37 | No | No | ˡYes | ᵐYes |
|  | L TP | -42, 4, -30 | 15 | 0.048 | 0.002 | 0.000 | 3.03 | ⁿYes | ᵒYes | ᵖYes | ʳYes |
| ᵃ ^a^Mean clusterwise functional connectivity from significant MVPA post-hoc seed-to-voxel clusters were entered into regression to predict residualized CAT-Q (partialing out age/SRS-2 variance) for each group. ˡF₍₇, ₁₃₎=6.50, p=.002, ᵐF₍₇, ₁₁₎=8.83, p=.001, ⁿF₍₃, ₂₀₎=5.71, p=.005, ᵒF₍₃, ₁₇₎=4.54, p=.016, ᵖF₍₃, ₁₇₎=4.00, p=.025, ʳF₍₃, ₁₅₎=3.26, p=.051 | | | | | | | | | | | |

| **Supplementary Table S2**. Post-hoc MVPA seed-to-voxel clusters surviving FDR-correction for “sex-typical,” “sex-atypical,” and “sex-agnostic” contrasts. | | | | | | | | | |
| --- | --- | --- | --- | --- | --- | --- | --- | --- | --- |
| Sex-atypical patterns | | |  |  |  |  |  |  |  |
|  | MVPA Seed | Connection | x, y, z | Size | Size p-FDR | Size p-unc | Peak p-unc | Peak p-unc | t_75_ |
|  | Precuneus | L MFG, SFG | -30, 16, 40 | 159 | 0.008 | 0.002 | <.0001 | <.0001 | -6.60 |
|  |  | R sLOC | 50, -70, 28 | 112 | 0.024 | 0.001 | <.0001 | <.0001 | -5.22 |
|  |  | L sLOC | -30, -76, 38 | 90 | 0.040 | 0.003 | <.0001 | <.0001 | -4.81 |
|  | Hypothalamus | Subcallosal; L ventral striatum, Forb, Thalamus, substantia nigra | -10, 12 -14 | 308 | <.0001 | <.0001 | <.0001 | <.0001 | 7.41 |
|  |  | R ACC | 0, 26, 24 | 97 | 0.042 | 0.001 | <.0001 | <.0001 | 5.06 |
|  |  | R/L cerebellum crus 2/1, R 6 | 8, -74, -30 | 174 | 0.003 | <.0001 | <.0001 | <.0001 | -6.02 |
| Sex-typical patterns | | |  |  |  |  |  |  |  |
|  | ACC | R/L PreCG, R/L PostCG | 26, -28, 68 | 821 | <.0001 | <.0001 | <.0001 | <.0001 | 6.42 |
|  |  | R IFG Oper/Tri, FO, Forb | 48, 20, -2 | 330 | <.0001 | <.0001 | <.0001 | <.0001 | 6.16 |
|  |  | R TP, aMTG | 50, 10, -28 | 146 | 0.006 | 0.000 | <.0001 | <.0001 | 6.01 |
|  |  | R pSMG, PostCG, SPL | 42, -36, 50 | 145 | 0.006 | 0.000 | <.0001 | <.0001 | 5.23 |
|  |  | L SPL, pSMG | -34, -42, 44 | 127 | 0.010 | 0.001 | <.0001 | <.0001 | 5.17 |
|  |  | ACC, PCC | -6, -12, 40 | 110 | 0.017 | 0.001 | <.0001 | <.0001 | 5.16 |
|  |  | L Thalamus | -8, -16, 8 | 94 | 0.028 | 0.002 | <.0001 | <.0001 | 4.85 |
|  |  | R Thalamus | 12, -20, 6 | 86 | 0.035 | 0.003 | <.0001 | <.0001 | 4.80 |
|  |  | L MFG, PreCG | -28, -4, 58 | 81 | 0.039 | 0.004 | <.0001 | <.0001 | 4.70 |
|  |  | R/L Lingual | -12, -84, -6 | 77 | 0.042 | 0.005 | <.0001 | <.0001 | 4.67 |
|  | R aPaHC | PCC, Precuneus | 6, -30, 38 | 254 | 0.000 | <.0001 | <.0001 | <.0001 | 6.10 |
|  |  | R SPL, AG | 32, -48, 42 | 154 | 0.005 | 0.000 | <.0001 | <.0001 | 5.77 |
|  |  | R AG, pSMG | 52, -54, 18 | 145 | 0.005 | 0.000 | <.0001 | <.0001 | 5.03 |
|  |  | L AG, sLOC | -52, -62, 26 | 116 | 0.013 | 0.001 | <.0001 | <.0001 | 4.98 |
|  |  | L Forb, IFG Tri | -52, 24, -8 | 108 | 0.014 | 0.001 | <.0001 | <.0001 | 4.96 |
|  |  | PCC | -10, -46, 28 | 88 | 0.029 | 0.003 | <.0001 | <.0001 | 4.67 |
|  |  | R TP | 38, 10, -50 | 134 | 0.014 | 0.000 | <.0001 | <.0001 | -5.81 |
| Sex-agnostic patterns (diagnosis differences) | | |  |  |  |  |  |  |  |
|  | R aPaHC | Precuneus; R sLOC, aSMG, pSMG, PostCG, AG | 40, -62, 46 | 1058 | <.0001 | <.0001 | <.0001 | <.0001 | 9.01 |
|  |  | R PreCG, IFG Oper, MFG | 54, 10, 8 | 375 | <.0001 | <.0001 | <.0001 | <.0001 | 7.33 |
|  |  | R FP, IFG Tri, MFG | 46, 30, 14 | 328 | <.0001 | <.0001 | <.0001 | <.0001 | 6.86 |
|  |  | L SPL, pSMG | -30, -52, 44 | 272 | <.0001 | <.0001 | <.0001 | <.0001 | 6.67 |
|  |  | PCC, Precuneus | 4, -44, 38 | 152 | 0.002 | 0.000 | <.0001 | <.0001 | 5.80 |
|  |  | R MFG, SFG | 34, 16, 58 | 112 | 0.008 | 0.001 | <.0001 | <.0001 | 5.61 |
|  |  | R MFG, FP | 32, 30, 44 | 100 | 0.012 | 0.001 | <.0001 | <.0001 | 5.39 |
|  | L TP | R AG, sLOC | 50, -60, 50 | 109 | 0.037 | 0.001 | <.0001 | <.0001 | 5.09 |
|  |  | Precuneus, L ICC, R Cuneus | 2, -70, 20 | 140 | 0.022 | 0.000 | <.0001 | <.0001 | -4.92 |
|  |  | L sLOC | -42, -70, 26 | 104 | 0.049 | 0.001 | <.0001 | <.0001 | -4.75 |
| *left (L), right (R), middle frontal gyrus (MFG), superior frontal gyrus (SFG), superolateral occipital cortex (sLOC), orbitofrontal cortex (FOrb), anterior cingulate cortex (ACC), precentral gyrus (PreCG), postcentral gyrus (PostCG), inferior frontal gyrus (IFG), temporal pole (TP), anterior middle temporal gyrus (aMTG), posterior supramarginal gyrus (pSMG), superior parietal lobule (SPL), angular gyrus (AG), posterior cingulate cortex (PCC), frontal pole (FP) | | | | | | | | | |

**Supplementary Figures**

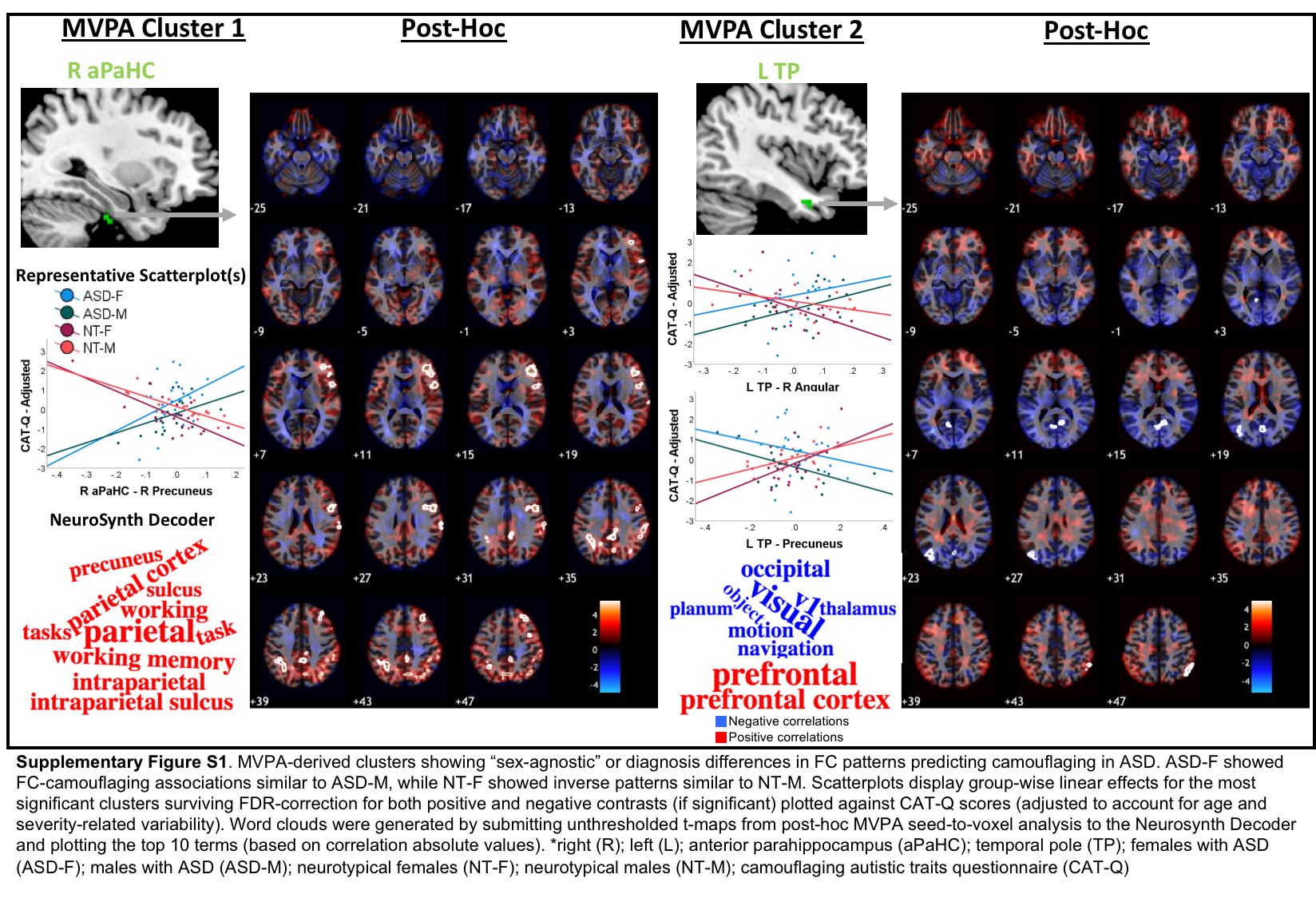

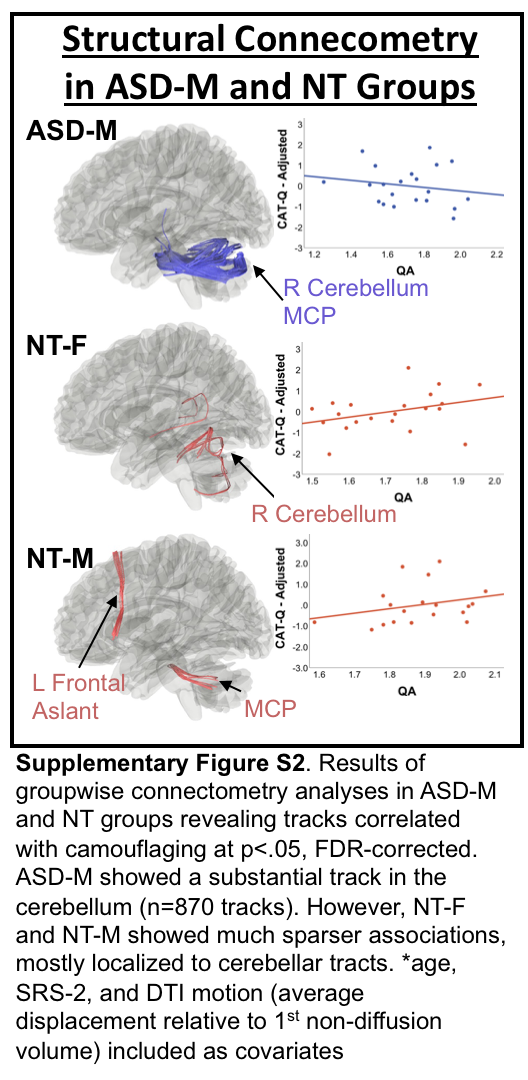

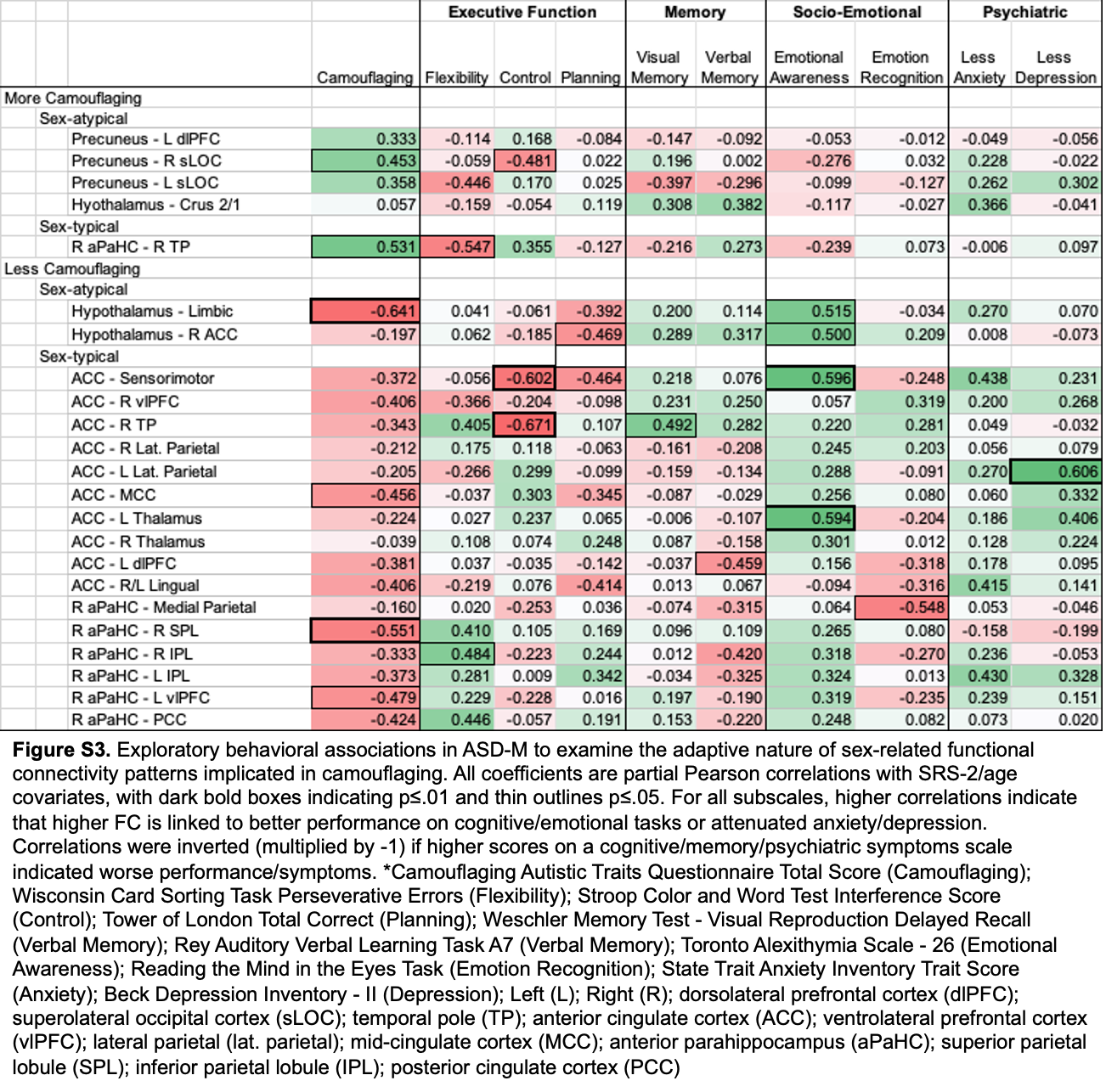

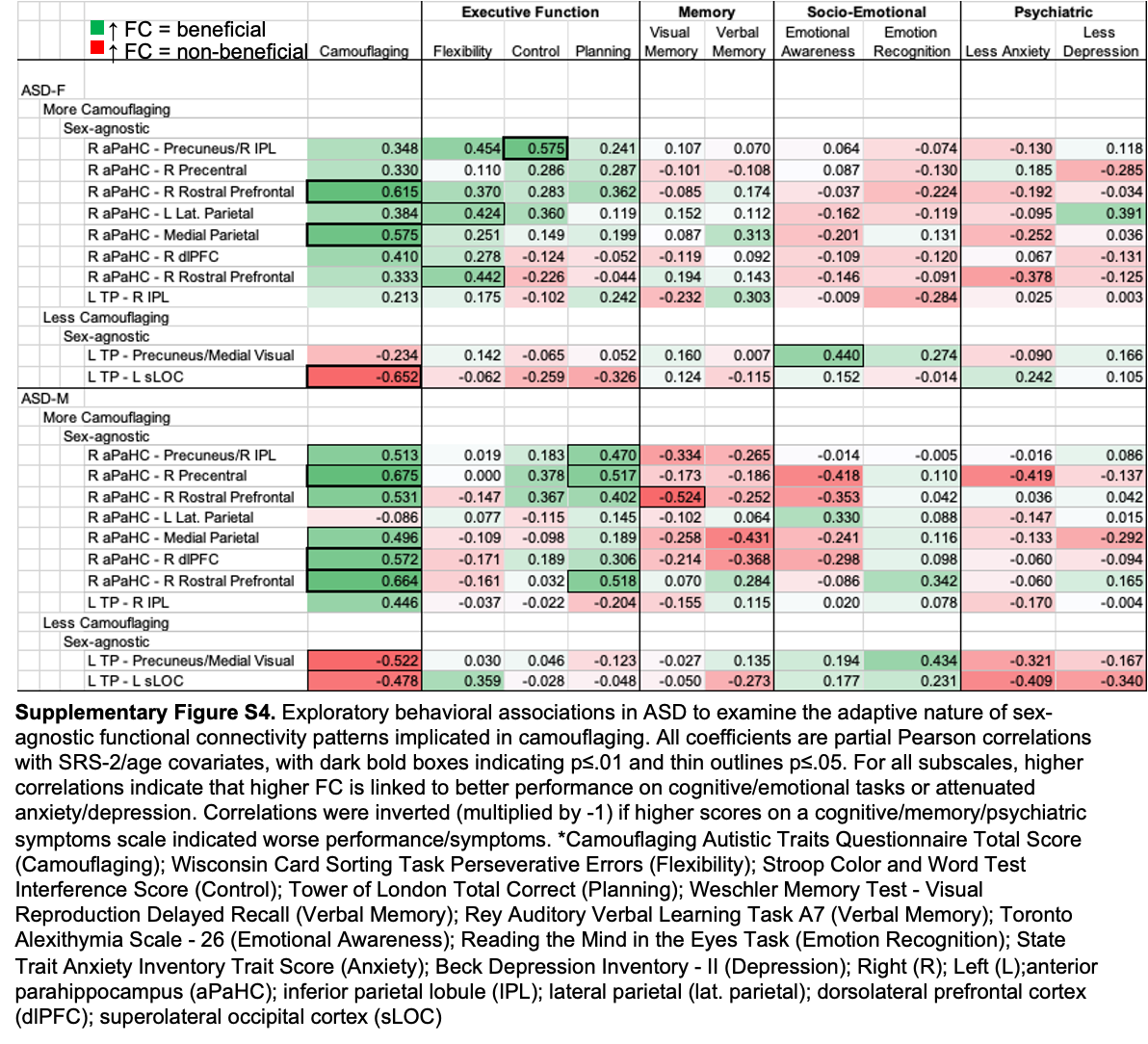
